# Microglial and Astrocyte priming in the APP/PS1 model of Alzheimer’s Disease: increased vulnerability to acute inflammation and cognitive deficits

**DOI:** 10.1101/344218

**Authors:** Ana Belen Lopez-Rodriguez, Edel Hennessy, Carol Murray, Anouchka Lewis, Niamh de Barra, Steven Fagan, Michael Rooney, Arshed Nazmi, Colm Cunningham

## Abstract

Alzheimer’s disease (AD) causes devastating cognitive decline and has no disease-modifying therapies. Neuroinflammation is a significant contributor to disease progression but its precise contribution remains unclear. An emerging literature indicates that secondary inflammatory insults including acute trauma and infection alter the trajectory of chronic neurodegenerative diseases and the roles of microglia and astrocytes require elucidation. The current study, using the APP/PS1 mouse model of AD, demonstrates that microglia are primed by β-amyloid pathology to induce exaggerated IL-1β responses to acute stimulation with LPS or IL-1β. Despite disease-associated NLRP3 inflammasome activation, evidenced by ASC speck formation, APP/PS1 microglial cells show neither IL-1β induction nor NFκB p65 nuclear localisation. Upon secondary stimulation with LPS or IL-1β, NFκB-p65 nuclear localisation and exaggerated pro-IL-1 induction occur. Microglial priming was also unmasked by secondary stimulation with systemic LPS leading to significant cognitive impairment in APP/PS1 mice compared to WT LPS-treated mice. Astrocytes have also recently emerged as displaying significant phenotypic heterogeneity. Here, by-passing microglial priming, and acutely challenging mice with intra-hippocampal IL-1β we demonstrate that astrocytes proximal to Aβ-plaques are also primed to produce exaggerated CCL2, CXCL1 and CXCL10 responses. Many astrocytosis-associated genes in APP/PS1 mice share these exaggerated responses to IL-1β, while others are equally induced in both strains. Collectively the data show that the amyloid-laden brain shows multiple vulnerabilities to secondary inflammatory challenge: both microglia and astrocytes are primed to produce exaggerated secondary inflammation and systemic LPS is sufficient to cause cognitive impairments relevant to delirium, selectively in animals with prior amyloid pathology.

## Introduction

Despite numerous clinical and preclinical studies, no disease modifying therapies for Alzheimer’s disease (AD), the top cause for disability in the elderly, have emerged. The majority of strategies used in prior studies have addressed amyloid or tau hypotheses (see (Umar and Hoda 2018) for review). Based on Genome wide association studies and extensive preclinical studies, neuroinflammation has emerged as an alternative target. Inflammation plays a contributory role in the development and progression of disease (Ardura-Fabregat et al. 2017) and this appears partly mediated through the response to secondary inflammation in or outside the central nervous system that exacerbates disease (Cunningham and Hennessy 2015).

Microglial and astrocyte populations are activated by neurodegenerative disease and detailed phenotypic analysis of these cells has been performed in recent years (Orre et al. 2014; Holtman et al. 2015; Keren-Shaul et al. 2017; Mrdjen et al. 2018) and has revealed robust activation of phagocytic machinery but relatively limited expression of NFκB-dependent genes and classical pro-inflammatory mediators such as IL-1β (Holtman et al. 2015). This suggests that the inflammatory profile of microglia during amyloidosis is, to some extent, restrained. Nonetheless, the NLRP3 inflammasome, an enzyme complex responsible for cleavage and maturation of IL-1β is activated and influences amyloidosis and cognitive function in the APP/PS1 model of AD (Heneka et al. 2012; Venegas et al. 2017). We previously showed that microglia are ‘primed’ by primary neuropathology to produce exaggerated acute neuroinflammatory responses to secondary acute stimulation with bacterial endotoxin (Cunningham et al. 2005), double stranded RNA (Field et al. 2010) and inflammatory cytokines IL-1β and TNFα (Hennessy et al. 2017). Such switching of glial phenotypes may release the restraint on microglia during neurodegeneration to induce a more classically activated microglial phenotype which may be an important component of the contribution of systemic inflammation to disease-associated outcomes such as neuronal damage and cognitive decline (Cunningham et al. 2009; Holmes et al. 2009; Semmler et al. 2013).

Although microglial priming was initially defined functionally, by this exaggerated IL-1 response to LPS (Cunningham et al. 2005), transcriptomic and weighted gene co-expression network analysis have now been used to define a molecular signature for primed microglia in multiple neurodegenerative models (Holtman et al. 2015). However, to our knowledge key ‘hub’ transcripts have not been demonstrated *in vivo* in animals with demonstrated exaggerated IL-1β responses to acute inflammatory stimulation. Moroever, evidence that microglia are primed in transgenic models of AD remains incomplete: using the Tg2576 murine model, exaggerated IL-1β mRNA was shown after i.p. LPS challenge but neither IL-1β protein nor a cellular source was established (Sly et al. 2001). Similarly, hypersensitive microglia were described in a presenilin-1 knock-in mouse (Mattson et al. 2002) but exaggerated *in vivo* microglial IL-1β synthesis was not demonstrated. Many studies have employed repeated systemic LPS dosing to study effects on Aβ, tau hyperphosphorylation, microgliosis and cognitive function (Sheng et al. 2003; Kitazawa et al. 2005; Lee et al. 2008; Bhaskar et al. 2010), but these have not explicitly examined whether microglia are primed by β-amyloid to produce exaggerated acute IL-1 responses. Finally, although extensive data sets suggesting significant microglial heterogeneity are emerging from single cell RNA sequencing studies (Damage associated microglia paper & Becher paper), these lack spatial information about localisation of different microglial phenotypes. Therefore the occurrence and spatial location of primed microglia in the amyloid-laden brain requires clarification.

Moreover, microglial activation is intimately integrated with astrocyte function. In AD models, there is evidence for astrocyte reactivity around Aβ deposits, showing hypertrophy, cytoskeletal changes and significant alterations in calcium activity, metabolism and innate immunity (Delekate et al. 2014). It has emerged that ‘reactive’ astrocytes, like microglia, can also adopt multiple phenotypes (Hamby et al. 2012; Zamanian et al. 2012; Hennessy et al. 2015; Liddelow et al. 2017; Clarke et al. 2018) and we have demonstrated, in ME7 mice with chronic neurodegeneration, that astrocytes also show primed responses: producing exaggerated chemokine (CCL2 and CXCL1) responses to acute stimulation with IL-1β or TNF-α (Hennessy et al. 2015). Here, we hypothesised that astrocytes in the APP/PS1 brain are primed to produce exaggerated chemokine responses to acute stimulation with IL-1β.

In the current work we aimed to 1) provide direct evidence for, and spatial resolution of, the priming of microglial cells, by amyloid pathology, to show exaggerated reponses to acute LPS stimulation, 2) reconcile this phenotype with recently published transcriptional signatures of microglial priming and with current understanding of NRLP3 inflammasome activation in APP/PS1 mice, 3) test the hypothesis that astrocytes are also primed in this model, to produce exaggerated chemokine responses to acute IL-1β stimulation and 4) investigate the vulnerability of these animals to acute cognitive impairment upon secondary inflammatory stimulation. The findings indicate multiple vulnerabilities of APP/PS1 mice to secondary inflammatory challenges and suggest an amplification loop involving exaggerated microglial responses as well as disproportionate responses of astrocytes to microglial secretory products.

## Materials and Methods

### Animals and stereotaxic surgery

APPSwe/PS1dE9 (Jax strain #005864, +/0) mice of 19±3 months were housed in cages of five or less at 21°C with a 12 h light/dark cycle. Food and water access was *ad libitum*. Mice were anesthetized i.p. with Avertin (50% w/v in tertiary amyl alcohol, diluted 1:40 in H_2_O; 20 ml/kg, i.p.; Sigma) and positioned in a stereotaxic frame (Kopf Instruments). Holes were drilled at 2.0 mm (anteroposterior) and 1.7 mm (right side of the midline) from Bregma, and 1 µl of LPS (1µg/µl) or IL-1β (10ng/µl) was injected into the right hippocampus using a glass microcapillary (Sigma) to a depth of 1.6 mm. The capillary was left *in situ* for 2 minutes before slow withdrawal. Control animals were administered 1 µl of saline 0.9%. After surgery, mice were placed in an incubator at 25°C for recovery. Animals were monitored for recovery from surgery. Further animals (17±1 months) were challenged i.p. with LPS (100µg/kg) before examination of acute sickness or assessment of cognitive function during acute systemic inflammation. All animal experimentation was performed under licenses granted by the Minister for Health and Children and from the Health Products Regulatory Authority, Ireland, with approval from the local ethical committee and in compliance with the Cruelty to Animals Act, 1876 and the European Community Directive, 86/609/EEC. Every effort was made to minimize stress to the animals.

### Acute sickness measurements

Temperature was measured at 0 h and 2 h after LPS challenge by a rectal probe (Thermalert TH5, Physitemp, Clifton, New Jersey), placed approximately 2 cm into the rectum of the mouse. Temperature deviation from baseline was calculated by subtracting the measurement at 0 h from the measurement at 2 h. To investigate locomotor and rearing activity the open field task was used. Mice were placed in a box (58×33×19 cm) and the number of times the mouse reared and crossed the squares in the box (distance travelled) was recorded for 3 min.

### Reference Memory and cognitive flexibility

To investigate reference memory and cognitive flexibility, the “paddling” Y-maze visuospatial task was used as described in (Cunningham et al. 2009). A clear perspex Y-maze consisting of three arms with dimensions 30 × 8 × 13 cm was mounted on a white plastic base. The distal end of each arm contained a hole, 4 cm in diameter and two arms could be blocked by insertion of a closed black plastic tube thus preventing mice from exiting. The third hole had an open black tube, 2 cm above the floor, where mice could exit the maze and enter a black burrowing tube to be returned to their homecage. Each of the three plastic tubes had a burrowing tube over them on the outside of the maze so that from the center of the maze all arms looked identical. The maze was filled with 2 cm of water at 20 to 22°C, sufficient to motivate mice to leave the maze by paddling to an exit tube,from where they are returned to their home cage. Mice were placed in one of two possible start arms in a pseudorandomised sequence for 10 trials and the groups were counter-balanced with respect to the location of the exit and start arm. For any individual mouse the exit arm was fixed (to test reference memory). WT and Tg mice (17±1 month) were trained for 6 days (10 trials per day). An arm entry was defined as entry of the whole body, excluding the tail and a correct trial was defined as entry to the exit tube without entering other arms. On day 7 mice were injected i.p with LPS (100ug/kg or vehicle) and at 2 hours post-challenge were tested on retention in the Y-maze for 1 trial to confirm retention of reference memory (based on pilot data indicating that retention was preserved under LPS challenge). The exit for each mouse was then switched to a different arm and mice had to learn the location of the new exit over 12 trials. The 12 trials were divided into 3 blocks of 4 and incorrect trials per block were recorded.

### Tissue preparation

Animals for analyses of cytokine-induced transcriptional changes were terminally anesthetized with sodium pentobarbital at 2 h post challenge (Euthatal; Merial Animal Health) and rapidly transcardially perfused with heparinized saline before dissection of hippocampus and snap freezing in liquid nitrogen and storage at -80°C until use.

Animals for immunohistochemical examination were terminally anesthetized with sodium pentobarbital (Euthatal; Merial Animal Health) and transcardially perfused with heparinized saline minutes followed by 10% neutral buffered formalin (Sigma). Brains were postfixed in formalin and then embedded in paraffin wax. Coronal sections (10 µm) were cut on a Leica RM2235 Rotary Microtome (Leica Microsystems) at the level of the hippocampus and floated onto electrostatically charged slides (Menzel-Glaser) and dried at 37°C overnight.

### RNA extraction

Total RNA was isolated using the RNeasy Plus Mini method (Qiagen, Limburg, Netherlands) following the manufacturer’s instructions. To ensure complete DNA elimination from the column bound RNA, an on-column DNase step was performed. The RNA yield and quality of each sample were quantified based on Optical Density (OD) using the NanoDrop’ND-1000 UV–vis spectrophotometer (Thermo Fisher Scientific). cDNA synthesis was carried out using a High Capacity cDNA Reverse Transcriptase Kit (Applied Biosystems, Warrington, UK). Primer and probe sets were designed using Applied Biosystems Primer Express software and amplified a single sequence of the correct amplicon size, as verified by SDS-PAGE. Where no probe sequence is shown, the DNA binding dye SYBR green was used in its place. Pre-made primers were also used and run following the manufacturer’s instructions (Thermo Fisher Scientific). Primer pair/probe sequences are shown in Table 1. Samples for RT-PCR were run in duplicate using FAM / SYBR (Roche) in a StepOne Real-Time PCR system (Applied Biosystems, Warrington, UK) under the cycling conditions: 95°C for 10 min followed by 95°C for 10 secs and 60°C for 30 secs for 40-45 cycles. Quantification was achieved by exploiting the relative quantitation method, using cDNA from LPS-injected mouse brain as a standard expressing all genes of interest and serial 1 in 4 dilutions of this cDNA to construct a linear standard curve relating cycle threshold (CT) values to relative concentrations, as previously described (Cunningham et al. 2005). Gene expression data were normalized to the housekeeping gene glyceraldehyde-3-phosphate dehydrogenase (GAPDH) and expressed relative to wild type saline-treated values.

**Table 1.**
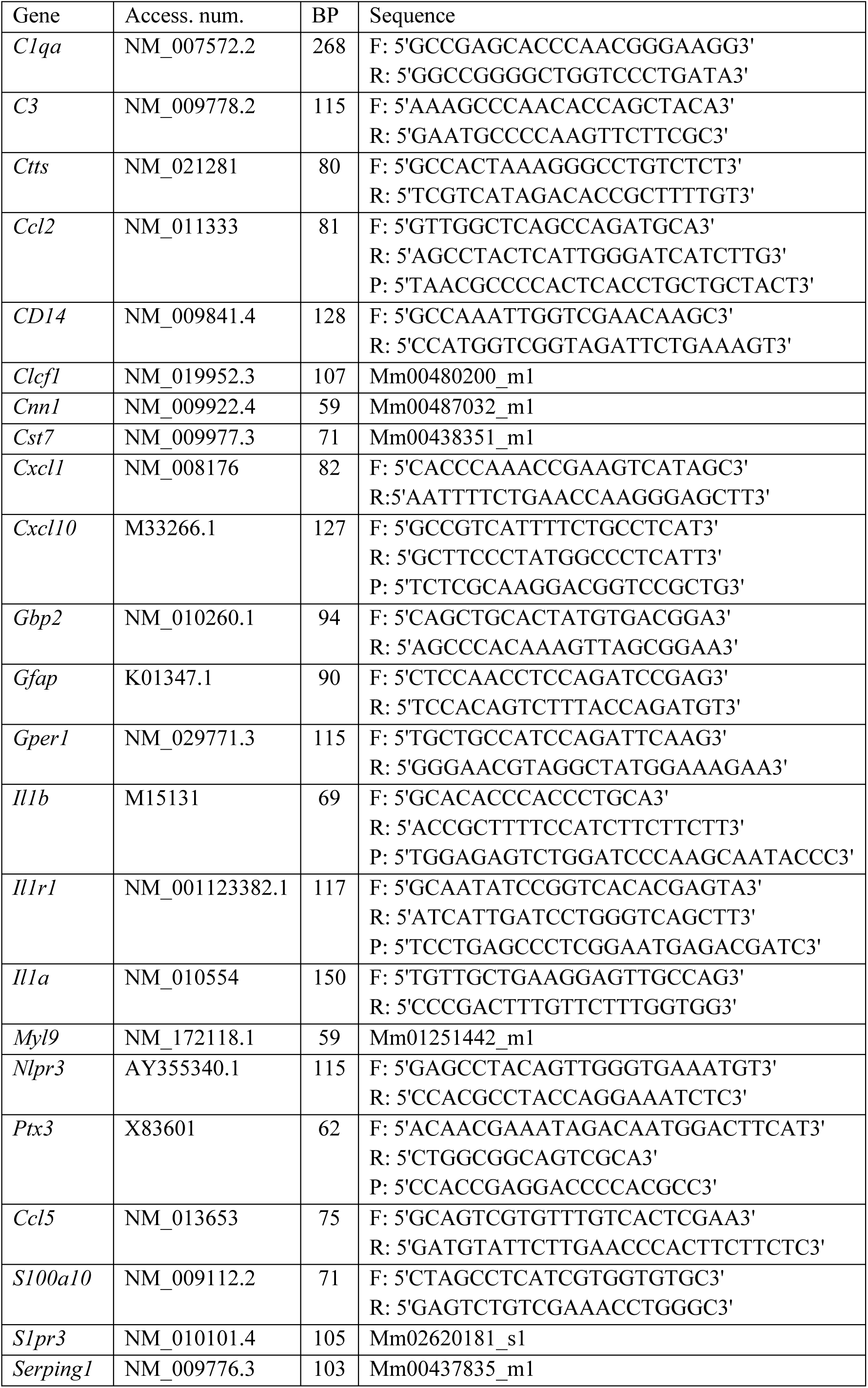
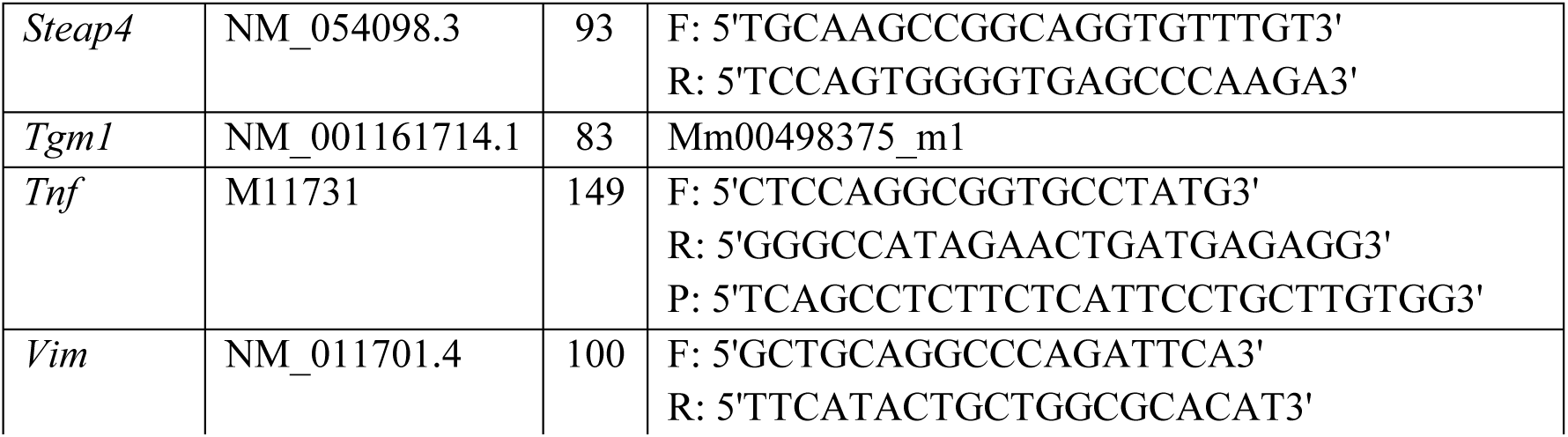
PCR *Primer sequences*. Where no probe sequence is shown, SYBR green dye was used. For pre-made primers, assay ID is provided. Forward (F), reverse (R), probe (P).

### Immunohistochemistry

Animals were labelled with GFAP (1:2000; Dako Z0334), 6E10 (1:1000; Biolegend 803001), Iba-1 (1:2000; Abcam ab5076), CCL2 (1:200; part 840288 of DY479 (R&D Systems), IL-1β (1:50; Peprotech 500-P51), CXCL1 (1:50; R&D AF-453-NA), CXCL10 (1:10000; Peprotech P129), NFκB (1:100; Santa Cruz sc-8008) and ASC (1:1000; AdipoGen AL177 006-C100). Briefly, all sections were quenched for 20 min with 1% H_2_O_2_/methanol and incubated overnight at 4 °C with the primary antibody and the appropriated serum. Next day, sections were incubated for 1 h with biotinylated secondary antibody (1:100, Vector). After several washes in phosphate buffer saline (PBS), ABC method was used (Vectastain Kit, PK6100 Vector) and the reaction product was revealed using 3, 3’ diaminobenzidine as chromogen (Sigma Aldrich) and H_2_O_2_ as substrate. Slides were counterstained using Haemotoxylin (VWR International Ltd, Dublin, Ireland), dehydrated, coverslipped, examined and photographed using an Olympus DP25 camera (Mason) mounted on a Leica DM3000 microscope (Laboratory Instruments and Supplies, Ashbourne), captured using CellA™ software (Olympus, Mason).

For confocal microscopy, sections were labelled with the following combinations: 6E10 (1:1000; Biolegend 803001) with an Alexa Fluor 488 anti-mouse (1:800; Invitrogen) together with Iba-1 (1:2000; Abcam ab5076) Alexa Fluor 594 anti-goat (1:800; Invitrogen); 6E10 with an Alexa Fluor 633 anti-mouse (1:800; Invitrogen) in combination with IL-1β (1:50; Peprotech 500-P51) with an Alexa Fluor 488 anti-rabbit (1:800; Invitrogen); 6E10 with an Alexa Fluor 633 anti-mouse and GFAP with an Alexa Fluor 488 anti-rabbit; GFAP (1:2000; Dako Z0334) with an Alexa Fluor 594 anti-rabbit (1:800; Invitrogen) together with IL-1β (1:50; Peprotech 500-P51) with an Alexa Fluor 488 anti-rabbit; GFAP with an Alexa Fluor 488 anti-rabbit in combination with CCL2 (1:200; R&D AF-479-NA) with Alexa Fluor 594 anti-goat; Iba-1 (1:2000; Abcam ab5076) with Alexa Fluor 594 anti-goat. The sections were incubated overnight at 4 °C with the primary antibodies and the appropriate serum. The day after, sections were incubated for 1 h with fluorescent secondary antibody and counterstained with Hoechst (1:2000; Sigma Aldrich 33258) for 10 min at RT. Slides were then coverslipped with ProLong™ Gold Antifade Mountant medium (Thermo Fisher Scientific) and examined with a Leica SP8 Scanning confocal microscope (Leica Microsystems).

### Mixed Glial Preparation, Isolation and Treatment

Cortices of C57BL6 mouse pups (p0-p4) were dissected out and dissociated using a Pasteur pipette and passed through a cell strainer (100 µm) into a 50 mL falcon. The cell suspension was added to T75 flask and incubated at 37°C in 15% DMEM (supplemented with 15% Horse Serum, 1% Glutamax and 1% penicillin/streptomycin; DMEM_supp_) with 10 ng/mL M-CSF and 5 ng/mL GM-CSF. At 90% cell confluency, microglia were isolated by centrifugation and seed onto a 24-well plate at a concentration of 6×10^4^ cells/well. 24h before treatment, media was changed and serum-free DMEM_supp_ was added. Cells were treated with IL-1β (2.5 ng/mL) for 6 h and the secretory profile of inflammatory mediators was measured by ELISA on the culture supernatant.

Astrocyte purification was done immediately after microglia isolation by cell separation columns kit (MACS®, Miltenyi Biotec). Remaining cells were resuspended in MACS beads / MACS buffer and run through the MACS columns. Purified astrocytes were seeded onto 6-well plates at a concentration of 2.5×10^5^ cells/well. 24h before treatment, media was changed to serum-free DMEM_supp_. Cells were treated with IL-1β (2.5 ng/mL) for 6 h and the secretory profile of inflammatory mediators was measured by ELISA on the culture supernatant, following manufacturers’ instructions.

### Statistical analyses

For multiple comparisons two-way analysis of variance (ANOVA) was performed, with factors being genotype (Wild Type or Transgenic) and treatment (LPS, Saline or IL1β). Data were not always normally distributed. Therefore, nonparametric tests were used (Kruskal–Wallis and post hoc pair-wise comparisons with Mann-Whitney U-test). Post hoc comparisons were performed with a level of significance set at p ≤ 0.05. For data that were normally distributed and homoscedastic, we used a standard parametric post-hoc test (Bonferroni’s test) and for those that were normally distributed, but non-homoscedastic, we performed non-parametric post-hoc comparisons (Games–Howell’s test). For two-group comparisons, data were analysed using Student-t test. Data are presented as mean ± standard error of the mean (SEM). Symbols in the graphs denote post-hoc tests. Statistical analyses were carried out with the SPSS 22.0 software package (SPSS, Inc., Chicago, IL, USA).

## Results

### Microglia are primed in APP/PS1 model

Microglia have been demonstrated to increase in number and reactivity around Aβ plaques (Glass et al. 2010; Mandrekar-Colucci and Landreth 2010; Wyss-Coray and Rogers 2012). Here we show, by immunohistochemistry, increased microglial activation in APP/PS1 mice (Fig.1A) and show a significant increase of the % area that is Iba-1-positive in Tg mice (Fig. 1B). The primed microglial gene signature was analysed by RT-PCR. *Sall1* maintains the down-regulated resident microglial phenotype (Buttgereit et al. 2016), while *Sparc*, *P2ry12* and *Tmem119* have roles in that maintenance and are reportedly decreased in isolated microglia populations of AD models (Keren-Shaul et al. 2017). We observed a significant decrease in *Sall1* (p=0.048) but no significant change in *Sparc*, *P2ry12* and *Tmem119*. However, there was a marked increase in *Tyrobp* (p<0.001), which reliably correlates with increased microglial numbers (Holtman et al. 2015; Wes, Holtman, et al. 2016; Wes, Sayed, et al. 2016), and significant increases in *Trem2*, *ITGAX* and *Clec7a* (p<0.001; Fig. 1C). These are microglial-specific genes and are part of the core gene signature for microglial priming (Holtman et al. 2015). These data suggest that these microglia have lost regulatory control and have become primed.

**Figure 1.**
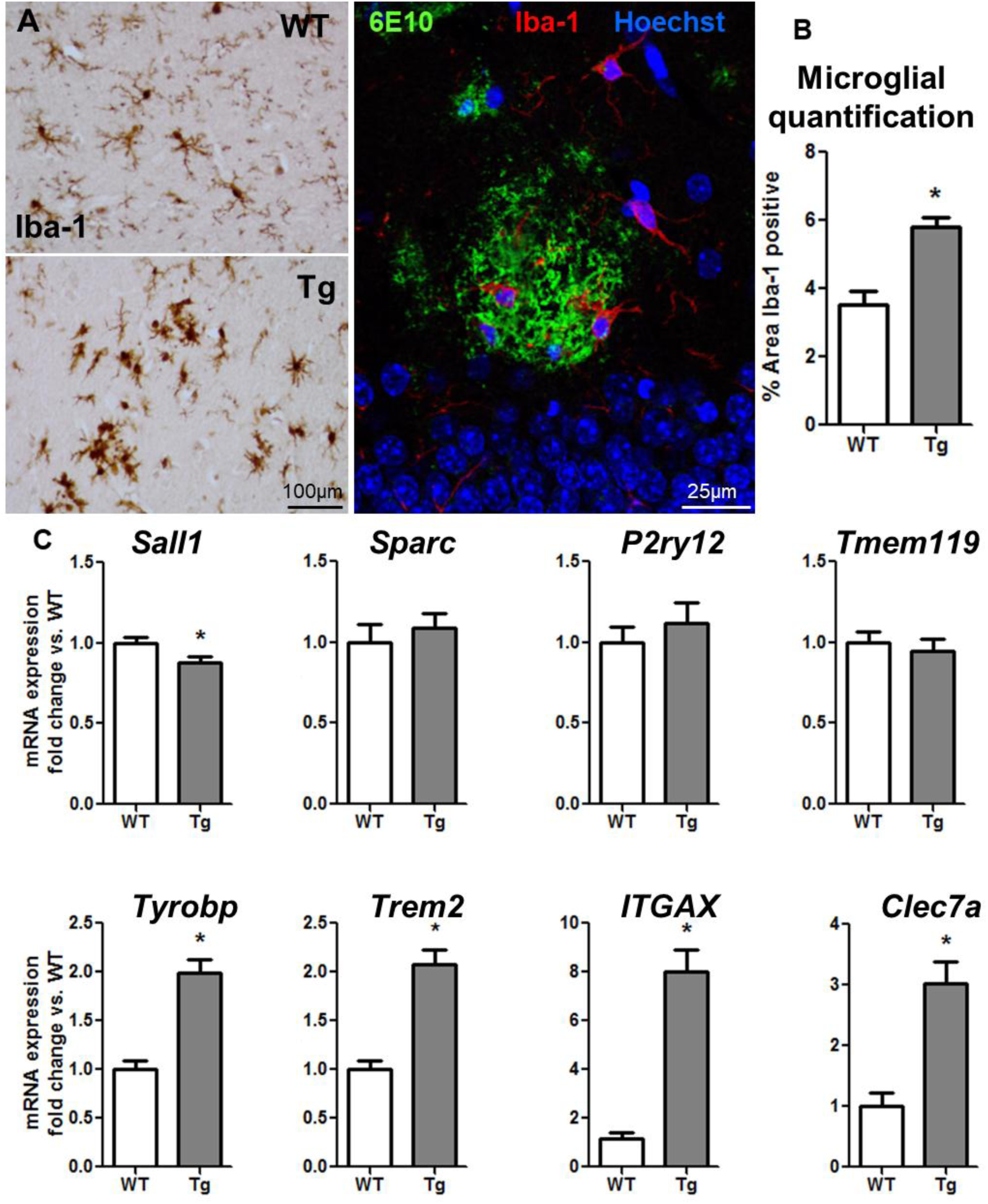
Microglial activation in APP/PS1. (A) Left panel: Comparison of microglial activation in WT and Tg (APP/PS1, 19±3 months) by Iba-1 labelling, light microscopy. Central panel: Confocal image of Tg brain tissue showing in (green, 488nm), Aβ plaque labelled with 6E10 surrounded by Iba-1 positive microglia (red, 594nm). (B) Microglial quantification in WT versus Tg represented as percentage of Iba-1-positive area. (C) mRNA expressed as fold change of specific microglial markers in WT and Tg (age 19±3months). Mean±SEM (WT n= 14; Tg n=19). t-test * vs. WT (p < 0.05).

Therefore we examined whether superimposed acute challenges (LPS or IL-1β i.c.), would produced exaggerated IL-1β responses in these microglia. Despite APP/PS1 mice showing a greater number of Iba-1 positive cells and more activated morphology than WT mice (Fig. 2A), IL-1β expression was not evident in Tg animals per se (2B, left). Acute stimulation with LPS induced exaggerated production of IL-1β in clusters of cells surrounding amyloid plaques in Tg animals (2B, centre) compared to isolated single IL-1β-positive cells in WT animals. This was also apparent in Tg mice challenged with IL-1β (Fig. 2B, right). This IL-1β is produced by microglial cells, specifically in those microglia directly adjacent to 6E10-positive Aβ plaques (Fig. 2C), but not by astrocytes, as shown by double-labelling with antibodies against IL-1β and IBA-1 or GFAP and confocal microscopy (Fig. 2C). This illustrates significant heterogeneity in microglial phenotypes within the amyloid-laden hippocampus: those adjacent to 6E10-positive plaques show exaggerated IL-1β response to local application of LPS.

**Figure 2.**
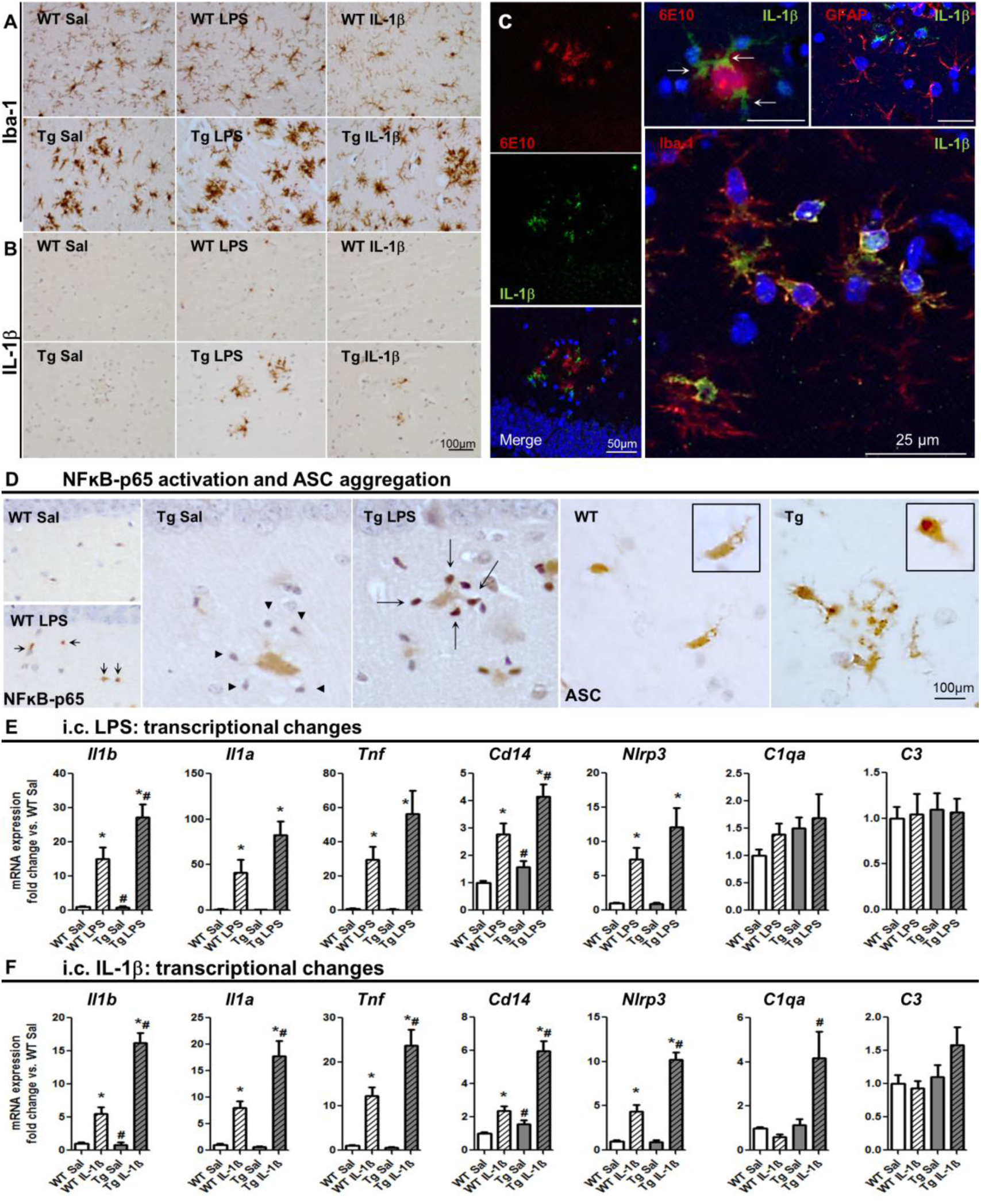
Microglial priming in APP/PS1 mice. Acute challenges (LPS or IL-1β i.c., 2h) superimposed on primed microglia in APP/PS1 (19±3 months). (A) Iba-1 labelling in WT and Tg treated with saline, 1 µg of LPS or 10 ng of IL-1β. (B) IL-1β labelling in the same animals. (C, left stack) Confocal micrographs from Tg+IL1β showing IL-1β positive cells (green, 488nm) around Aβ plaque stained with 6E10 in (red, 630nm). Larger panel shows IL-1β positive cells (green, 488nm) co-localised with Iba-1 positive cells (red, 594nm). Upper right panel shows IL-1β positive cells (green, 488nm) that do not overlap with red GFAP-positive cells (594nm). (D) NFkB-p65 labelling (left panels) showing lack of activation in Tg+sal nuclei (arrowheads) and activation in Tg+LPS (arrows). ASC aggregation (right panels) showing speck formation in Tg but not WT animals (zoom in the insets). (E) mRNA expressed as fold change of specific microglial markers after LPS i.c. (2h). (F) mRNA expressed as fold change of selective microglial markers after IL-1β i.c. (2h). Mean ±SEM (n=8-10). Kruskal Wallis Test. * effect of treatment; # effect of genotype (p< 0.05).

To better understand this phenotypic switch in plaque-associated microglia we assessed two key steps in the synthesis and maturation of IL-1β. The nuclear translocation of the p65 subunit of nuclear factor kappa-light-chain-enhancer of activated B cells (NFκB) is an essential step in initiation of the expression of *Il1b* and *Nlp3*, a key constituent of the NLRP3 inflammasome complex (often referred to as signal 1). Thereafter, when NLRP3, ASC and pro-caspase 1 are assembled, the active form of caspase-1 cleaves pro-IL1β to induce its maturation and secretion (Latz et al. 2013; White et al. 2017). Microglia adjacent to Aβ plaques in the hippocampus do not show nuclear localisation of NFkB-p65 in Tg mice challenged only with saline, with nuclei surrounding plaques showing only blue haematoxylin counterstain (arrowheads in Fig. 2D). In contrast, the Tg+LPS group show clear nuclear translocation of NFkB-p65 in the cells surrounding plaques (arrows Fig. 2D). This conclusion is supported by minimal expression of either *Il1b* or *Nlrp3* in Tg+Sal animals but distinct and indeed exaggerated induction of these genes in Tg+LPS compared to both Tg+Sal and WT+LPS groups (Fig. 2E).

Examining inflammasone activation using ASC aggregation (often referred to as signal 2), WT animals showed modest expression and uniform distribution of ASC in microglia of the hippocampus while in Tg animals (untreated) we observed more intense labelling in plaque-associated microglia (Fig. 2D, right), with a non-uniformity of ASC distribution within the cell consistent with ASC “speck” formation within these cells (insets). Therefore, plaque-associated microglia in Tg animals show inflammasome activation (signal 2) as a result of amyloidosis, as was previously shown (Heneka et al. 2012; Venegas et al. 2017), however we now show that these cells do not translocate NFκB p65 to the nucleus and do not present significant pro-IL-1β to the assembled inflammasome (i.e. no signal 1). However, these microglia are primed to transcribe and translate exaggerated IL-1β upon secondary stimulation and are equipped to process this IL-1β effectively.

RT-PCR analysis of cytokine, inflammasome and complement genes 2 hours after an acute LPS challenge (i.c.) showed that LPS-treated animals of both genotypes showed an increase in *Il1b*, *Il1a*, *Tnf*, *CD14* and *Nlrp3* mRNA levels, with no changes in *C1qa* or *C3*. The effects of LPS (*) and genotype (#) on expression were assessed using Kruskal Wallis test, followed by Mann-Whitney analyses. Within this set of genes, *Il1b* and *CD14* presented an exaggerated response in Tg+LPS in comparison with WT+LPS (Fig. 2E; all p values <0.043).

Since LPS rarely gains access to the brain, we also assessed whether microglia would show exaggerated responses to IL-1β, since this may arise in multiple traumatic and infectious episodes. We directly injected IL-1β i.c. and showed that only microglia in the Tg brain showed detectable IL-1β protein expression at 2 hours (Fig. 2B). Examining transcriptional changes 2 hours after IL-1β challenge (Fig. 2F), we showed that IL-1β-induced increases in *Il1b*, *Il1a*, *Tnf*, *CD14* and *Nlrp3* mRNA levels but exaggerated increases in *Il1b*, *Il1a*, *Tnf*, *CD14*, *Nlrp3* and *C1qa* mRNA expression in Tg+IL-1β animals compared to WT+LPS (all p values < 0.041, effect of IL-1β (*) or genotype (#)).

### Systemic inflammation also unmasks microglia priming and causes cognitive impairment

Having demonstrated that APP/PS1 microglia were primed to produce exaggerated responses to central LPS/IL-1β challenge, we also assessed whether this exaggerated response could also be triggered by systemic inflammation (i.p.). We used *CD68* and *C1qa* mRNA levels (Fig. 3A) to show that this discrete cohort of animals showed the expected increased complement and phagocytic activation known to occur with amyloidosis (Hong et al. 2016). Examining a similar panel to those examined after i.c. LPS we found a significant effect of systemic LPS in WT and Tg mice in *Il1b*, *Tnf* and *CD14* and exaggerated *Il1b* and *CD14* responses in the Tg+LPS group (all p values < 0.047). The analysis of functional consequences of such challenges showed that systemic LPS (100 µg/kg) induced sickness behaviour in both strains as indicated in Fig. 3 C,D. Animals challenged with LPS showed a decrease in body temperature and in distance travelled in the open field but there were no differences in those effects between WT and Tg animals, indicating equivalent sickness. When animals were tested on retention of previously learned visuospatially-led exit from the Y-maze, LPS did not impair this function (data not shown). However, when the location of the exit was moved to an alternate arm, thus requiring cognitive flexibility and the adoption of a new strategy for maze escape, we observed significant impairment in Tg+LPS animals with respect to all other animals (Fig. 3E). Kruskal Wallis Test revealed a significant effect of genotype (#) and i.p. LPS (*) and significantly worse cognitive impairment in Tg+LPS (all p values < 0.032).

**Figure 3.**
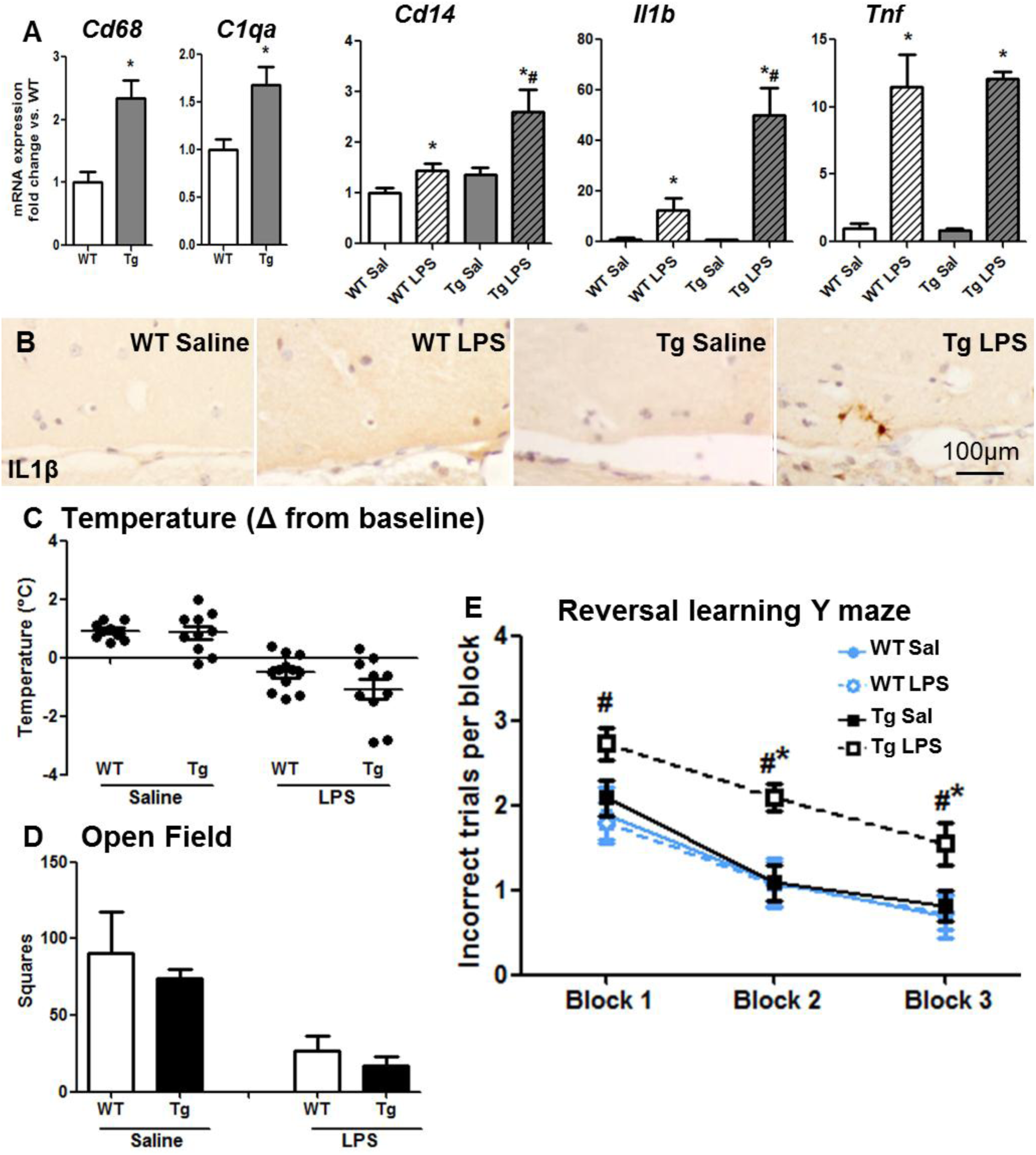
Systemic LPS produces exaggerated microglial responses and acute cognitive dysfunction. (A) mRNA expressed as fold change of selective microglial markers in APP/PS1 and WT mice (17±1 month) 2h after LPS (100 µg/kg i.p.) challenge. (B) IL-1β staining in animals under the same conditions. Mean ±SEM (n=5-7). (C) Deviation from baseline core-body temperature. (D) Locomotor activity in 3 minutes in the open field test. (E) Reversal learning Y maze. Mean ±SEM (n=10-15). Kruskal Wallis Test. * vs. WT in t-tests; * effect of treatment; # effect of genotype (p < 0.05).

### Astrocytes are primed in APP/PS1 mice

Astrocytosis is a well-described feature of AD and relevant mouse models (Ben Haim et al. 2015). Despite long-standing descriptions of astrocytes simply as ‘reactive’ there is an increasing awareness that there is significant heterogeneity in astrocyte phenotypes and these include clear evidence of innate immune activation (Zamanian et al. 2012; Orre et al. 2014; Ben Haim et al. 2015; Liddelow et al. 2017). We have previously described that astrocytes are primed during neurodegeneration in ME7 prion disease to show exaggerated chemokine expression in response to locally applied IL-1β or TNF-α (Hennessy et al. 2015) and here we investigated the hypothesis that similar ‘priming’ occurs in the APP/PS1 model of AD. We analysed a panel of genes identified by previous investigators as being elevated in astrocytes in models of stroke, sepsis and AD (Zamanian et al. 2012; Orre et al. 2014; Liddelow and Barres 2016) while also attending to recently described A1, A2 and pan reactive signatures (Liddelow et al. 2017).

Immunohistochemistry for GFAP shows astrogliosis encircling a 6E10-positive Aβ plaque in Tg mice (Fig. 4A left) and this is more evident using confocal microscopy showing GFAP-positive astrocytes (green) surrounding the 6E10-labelled plaque (red). The transcriptional signature was categorised into three groups: those transcripts affected by disease (Fig. 4B), those altered by acute IL-1β challenge (Fig. 4C) and those showing exaggerated responses to IL-1β (Fig. 4D). *Gfap*, *Ctss*, *Serping1* and *Il1r1* showed a significant increase in disease (all p values < 0.036) but were not affected by IL-1β i.c. challenge. Acute IL-1β produced a significant increase in *Steap4*, *Ccl2*, *Cxcl1*, *Ccl5* and a decrease of *Gper1* mRNA expression (Fig. 4C; all p values < 0.018) but no significant disease or IL-1-mediated increase in *Vim*.

**Figure 4.**
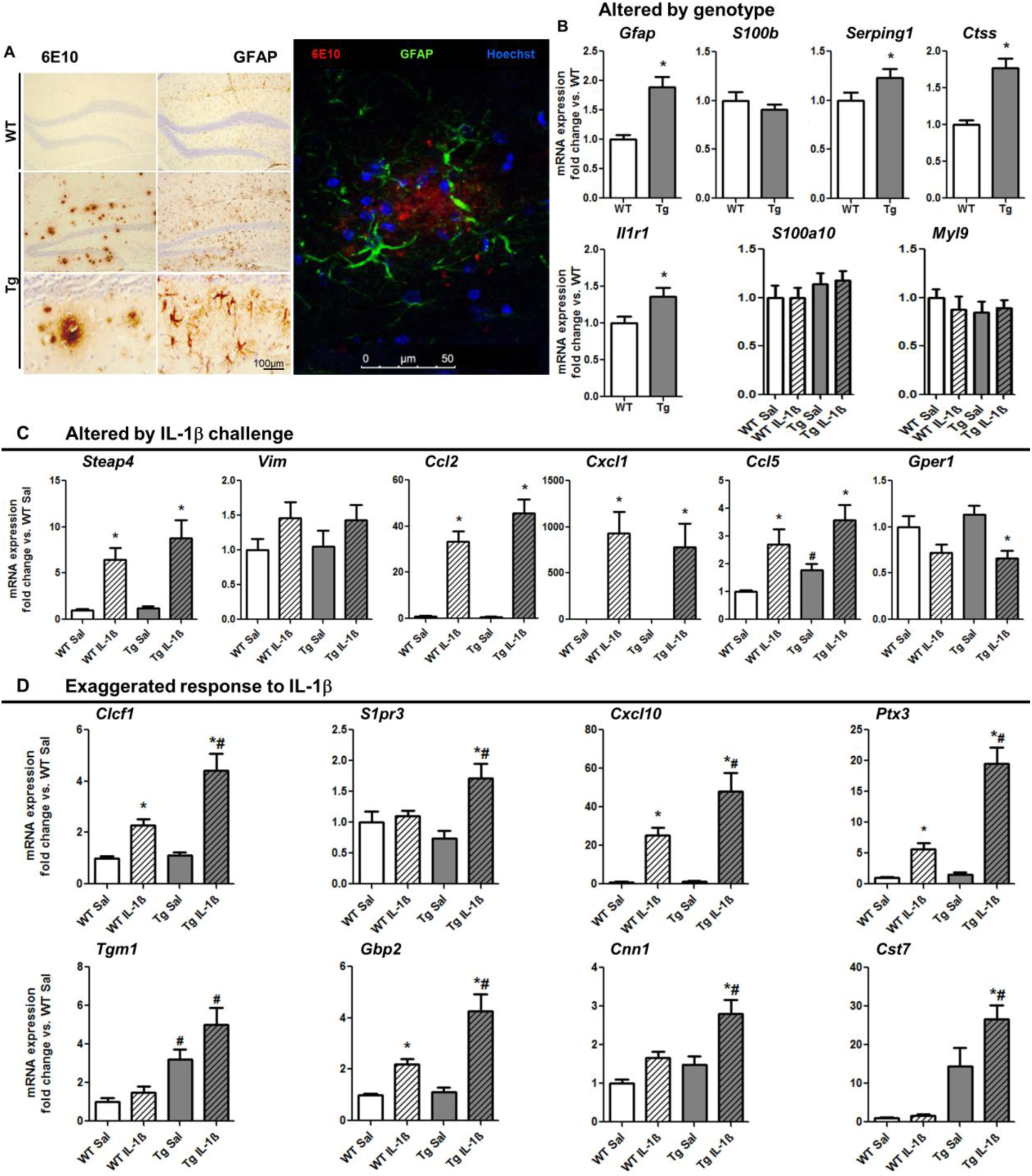
Astrocyte activation in APP/PS1 ± IL-1β (10 ng) i.c. challenge (2h). (A) Light microscopy labelling of 6E10 (left) and GFAP (right) of WT (first line) and Tg (second and third line). On the right panel, confocal image of Tg brain tissue labelled with 6E10 in red (630nm), green GFAP-positive astrocytes (488nm) surrounding the plaque. Transcriptional changes (mRNA) in selected astrocytic genes, expressed as fold change with respect to WT+saline: (B) altered exclusively by genotype, (C) altered by i.c. IL-1β challenge and (D) showing an exaggerated response in Tg animals treated with IL-1β. Aged 19±3months. Mean ±SEM (n=8-10). * vs. WT in T-test; Kruskal Wallis Test, * effect of treatment; # effect of genotype (p < 0.050).

Several transcripts showed differential responses to IL-1β challenge in Tg and WT animals (Fig. 4D). Although IL-1β induced several further transcripts in WT mice (*Clcf1*, *Cxcl10*, *Ptx3* and *Gbp2*; all p values <0.030) and many were increased in disease per se (*Clcf1*, *S1p3*, *Cxcl10*, *Ptx3*, *Gbp2*, *Cnn1* and *Cst7*; all p values < 0.027) several genes were induced to an exaggerated extent by IL-1β in Tg mice. Using Kruskal Wallis Test followed by Mann-Withney analyses, we found an effect of genotype and treatment exclusively in Tg+IL-1β group, meaning an interaction between treatment and genotype for all transcripts in Fig. 4D (all p values < 0.035).

These exaggerated responses of known astrocytosis-associated transcripts to acute IL-1β treatment are indicative of astrocyte priming but to directly address the hypothesis that the astrocytes are primed we focussed on chemokine expression. It is known that both astrocytes and microglia are capable of chemokine synthesis. We used *in vitro* cultures of primary glial populations to show that astrocytes synthesise greater levels of the chemokines CCL2, CXCL1 and CXCL10 than do microglia, in response to IL-1β (2.5 ng/mL for 6h). Although microglial cells showed higher basal release of CCL2, the treatment of cultured microglia with IL-1β revealed no significant increase in the release of CXCL1, CCL2 or CXCL10 (Fig. 5A-C). Conversely astrocytes showed clear CXCL1 (p<0.0001) and CCL2 (p<0.05) expression in response to IL-1β treatment and a trend towards increased CXCL10 release (p=0.057). Thus primary astrocytes produce more chemokines than microglia under IL-1β stimulation. In our prior studies in the degenerating hippocampus of mice with the ME7 strain of prion disease, exaggerated CCL2 and CXCL1 expression in response to acute cytokine stimulation proved to be a robust measure of the primed status of astrocytes during evolving pathology (Hennessy et al. 2015). Therefore, here we examined chemokine production in APP/PS1 mice using immunohistochemistry (Fig 5. D-G). Since these chemokines are also robustly expressed at the brain endothelium upon acute IL-1β stimulation this served as a positive control for the immunolabelling method.

**Figure 5.**
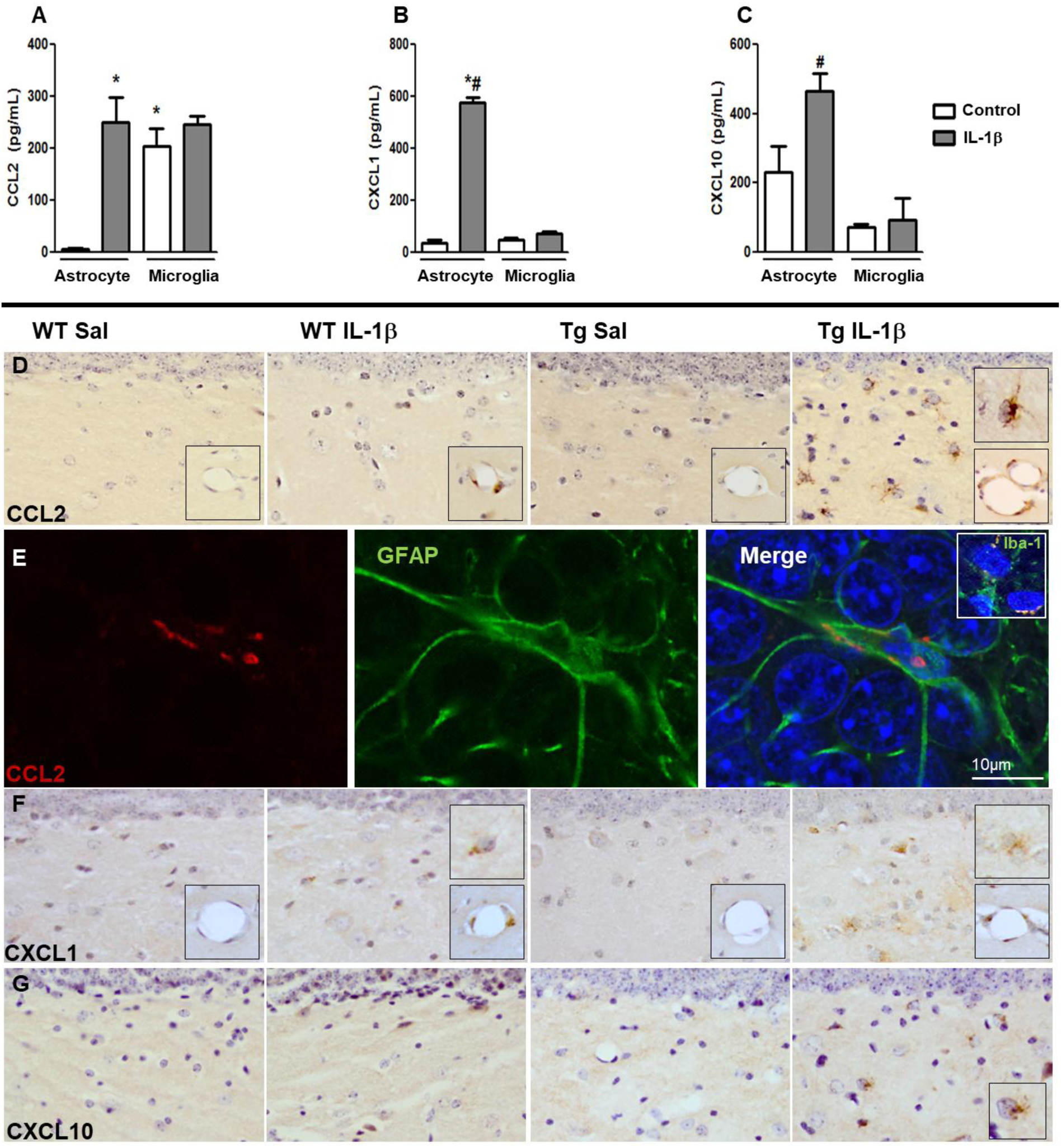
Astrocytes are primed to show exaggerated chemokine production in APP/PS1. (A-C) Chemokine secretion in microglia and astrocyte supernatants, 6h after IL-1β challenge (2.5 ng/mL) measured by ELISA for CCL2, CXCL1 and CXCL10. Two way ANOVA, *p<0.001, compared to astrocyte control; #p<0.001, compared to microglia IL-1β. (D) Light microscopy labelling of CCL2 in WT and Tg mice, 2h after i.c. challenge with IL-1β (10 ng) or saline. (E) Confocal imaging of Tg+IL-1β group for CCL2 (red; 594nm) and GFAP (green; 488nm). Iba-1 (green; 488nm) is captured in the inset. (F) Light microscopy labelling of CXCL1. (G) Light microscopy labelling of CXCL10. Insets show detail of positive vascular and astrocytic cells for the different chemokines.

Although CCL2 labelling was clear in the vasculature of both WT and Tg animals challenged with IL-1β, only Tg+IL-1β animals showed clear labelling of parenchymal cells. CCL2-positive cells in the dentate gyrus had large circular nuclei typical of astrocytes and clearly distinguishable from the smaller, darker microglial nuclei (Fig. 5D right panel). The identity of CCL2-positive cells as astrocytes was confirmed by confocal imaging (Fig. 5E), demonstrating perinuclear punctate CCL2 labelling (red) in astrocytes (green). Some CCL2 labelling was also observed in microglia (green) but this tended to present as a single perinuclear puncta (inset).

Similarly CXCL1 expression was clear in the vasculature of both IL-1β-treated groups while only Tg+IL-1β animals showed significant parenchymal expression of CXCL1 and this was once again in cells showing astrocyte morphology (Fig. 5F and inset). Likewise CXCL10 immunoreactivity was only found in Tg animals challenged with IL-1β (Fig. 5G), showing astrocyte morphology (inset). Thus astrocytes proximal to Aβ-plaques in APP/PS1 mice are primed to show exaggerated chemokine responses to acute IL-1β stimulation.

## Discussion

The present study shows that both microglia and astrocytes are primed in the APP/PS1 model. While microglia show an exaggerated IL-1β response to LPS stimulation, astrocytes show an exaggerated chemokine response to IL-1β, suggesting an amplification loop that drives exaggerated inflammatory responses to acute stimulation in the vulnerable AD brain (Fig. 6). Consistent with negative consequences of these vulnerabilities to secondary inflammatory stimulation, APP/PS1 mice were more vulnerable to acute cognitive dysfunction triggered by bacterial LPS.

**Figure 6.**
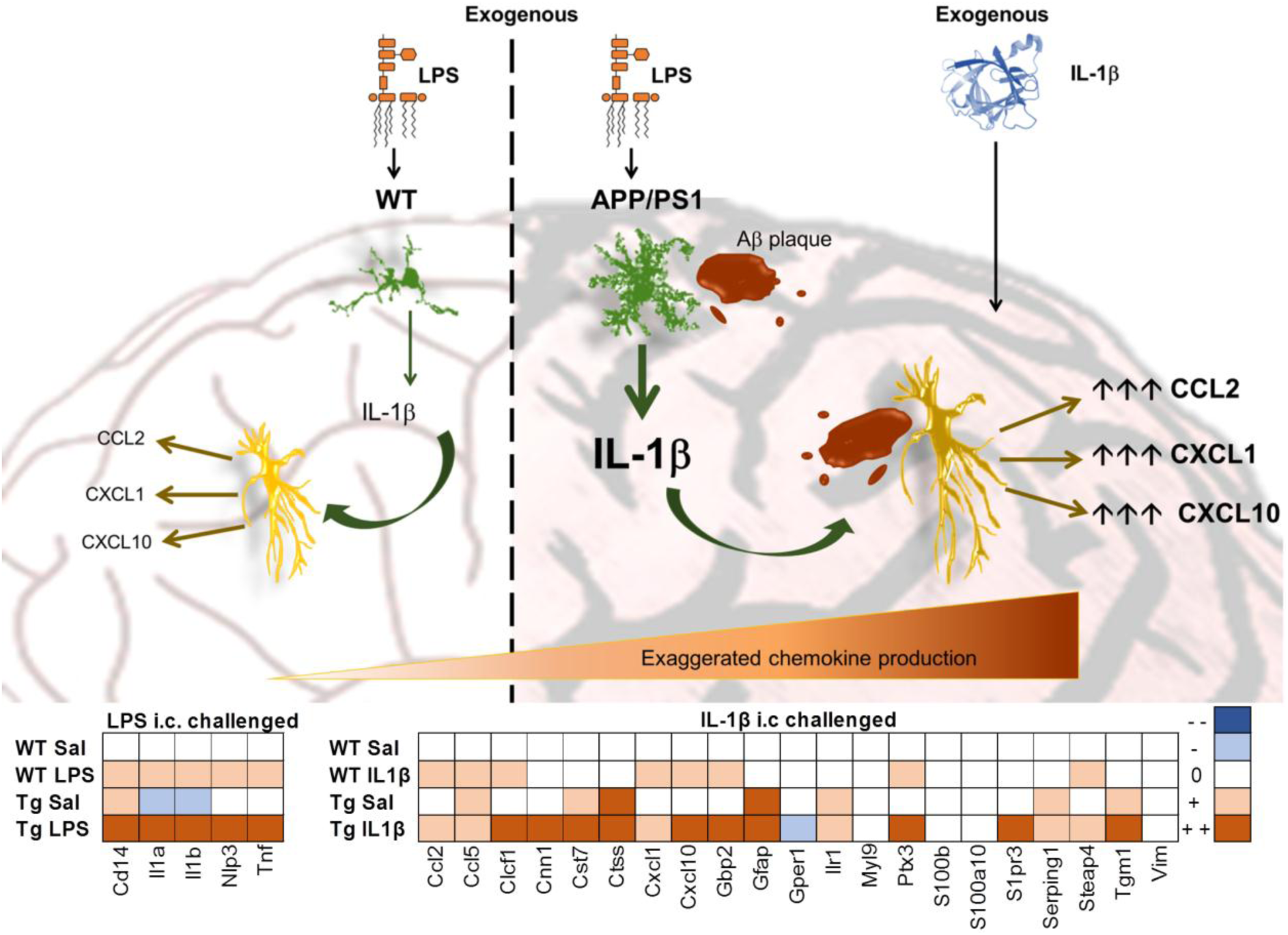
Microglial and astrocyte priming in APP/PS1 drives an inflammatory amplification loop. Exogenous i.c. administration of LPS in normal mice induces activation of microglial cells to produce regulated secretion of IL-1β and this IL-1β may activate astrocytes to release chemokines such as CCL2, CXCL1 or CXCL10. However, in APP/PS1 mice, LPS stimulates more robust production and processing of IL-1β from microglia primed by Amyloid pathology. Astrocytes primed by amyloid pathology also respond in an exaggerated way to exogenous IL-1β, producing enhanced levels of chemokines. If levels of IL-1, arising endogenously from microglial stimulation, are already exaggerated this will produce a further amplification upon stimulating astrocytes that are hypersensitive to IL-1. These exaggerated responses also appear to affect many other astrocytic transcripts involved in innate immune function as shown by fold-change expression in comparison to WT+saline.

### Microglial priming

We sought, in the APP/PS1 model, to provide direct evidence reconciling the recently emerged detailed transcriptomic signature for microglial priming (Holtman et al. 2015) with the functional definition of primed microglia i.e. that microglia show enhanced IL-1β responses to secondary stimulation (Cunningham 2013). As previously shown, the microglia of APP/PS1 mice (19 months) showed increased Iba-1-positive area compared to the healthy brain, with thickening and shortening of the processes, selectively proximal to Aβ plaques. Holtman et al., described elevation of ‘hub’ genes of microglial priming (*Clec7a, Itgax*), elevation of microglial genes strongly correlating with microglial quantity (*TyroBP, Trem2*) and suppression of expression of what are generally termed homeostatic genes such as *Sall1* (Buttgereit et al. 2016), *Sparc* (Lloyd-Burton et al. 2013), *P2ry12* (Haynes et al. 2006) and *Tmem119* (Satoh et al. 2016). Here we confirmed increases in *Tyrobp*, *Trem2*, *Itgax* and *Clec7a* and a significant decrease in *Sall1*, a key regulator for maintenance of the surveilling, homeostatic microglial phenotype (Buttgereit et al. 2016). Although the homeostatic transcripts *Sparc*, *P2ry12* and *Tmem119* were not significantly reduced, their failure to be upregulated despite the increased microglial numbers probably reflects a decreased level of expression of these transcripts per cell. Collectively the data are consistent with the previously reported transcriptional signature of primed microglia in the APP/PS1 model (Holtman et al. 2015). Functionally, microglia proximal to Aβ produced exaggerated IL-1β responses to acute stimulation with LPS and indeed to stimulation with IL-1β. Therefore multiple stimuli can evoke an exaggerated response from microglia if they have been primed by the prior stimulus, in this case Aβ plaques. *De novo* IL-1β was synthesised by activated microglia, but not by astrocytes, as corroborated by the confocal imaging

Microglia throughout the hippocampus did not produce IL-1β after i.c. challenge with LPS. Rather those microglia surrounding the plaques produced intense labelling with antibodies against IL-1β. Although recent data showing significant microglial heterogeneity within the AD transgenic brain were generated using an elegant single cell RNAseq approach (Keren-Shaul et al. 2017; Mrdjen et al. 2018), those studies cannot show the precise spatial resolution of these phenotypes. Here we clearly show, using classical immunohistochemical approaches, that those cells that are primed and that show exaggerated IL-1β responses to secondary stimulation are specifically those directly adjacent to amyloid plaques.

### Inflammasome activation: contribution to microglial priming

In order for IL-1β to be released it must be transcribed, translated, cleaved and secreted and processing of IL-1β is facilitated by the formation of the inflammasome (Rathinam et al. 2012). Following an appropiate inflammatory stimulus, NFκB p65 translocates to the nucleus to activate the transcription of pro-IL-1β and *Nlrp3* (often termed signal I, but by unfortunate coincidence, also called “priming” of the inflammasome). The assembly of the inflammasome complex from NLRP3, ASC and caspase 1 is typically induced by an additional stimulus (signal II or “activation”) (Hornung and Latz 2010; Lopez-Castejon and Brough 2011; Latz et al. 2013), and there is good evidence that Aβ triggers this in models of Alzheimer’s disease (Halle et al. 2008; Semmler et al. 2013; Venegas et al. 2017). Our data are consistent with this: ASC assembles into a large protein “speck” and this change from uniform distribution in the cytoplasm to the clustered or aggregated from is a key descriptor of inflammasone activation (Halle et al. 2008). Here we show uniform staining of ASC in microglia of WT animals and clear speck formation in microglia around plaques of Tg animals. However, the strength of the pro-IL-1 signal is weak. We found no evidence of increased *Il1b* mRNA in APP/PS1 *per se* and there was no nuclear localistion of NFkB p65 in nuclei of microglia surrounding the plaque. In earlier studies microglia were first “primed” with LPS in order to provide a pro-IL-1 signal for the assembled inflammasome to process (Halle et al. 2008) and the data here support the idea that, in the APP/PS1 brain, the microglial NLRP3 inflammasome is assembled during disease and has the capacity to cleave pro-IL-1 but there is a weak supply of pro-IL-1β to cleave. However, under secondary inflammatory stimulation, with LPS or IL-1β, microglia now amply synthesise pro-IL-1β and this can then be rapidly processed. A surprising implication of these data is that the classically described signal I, in the current model, is occurring *after* signal II. We do not rule out that *Il1b* synthesis and *Nlrp3* mRNA were transcribed at some earlier point in disease and are simply no longer being transcribed at this later stage. However it appears that, with the inflammasome assembled, later ‘pulses’ of pro-IL-1β arising from secondary inflammation, which are exaggerated due to the priming of microglia, can readily provide further IL-1 for processing, which may have significant consequences for cognitive function and pathology.

### Astrocyte Priming

Microglial responses synergise with astrocyte activation to shape, and to respond to, the extracellular milieu. Here we demonstrate that astrocytes are primed in the APP/PS1 model to produce exaggerated chemokine responses to acute IL-1β stimulation. Although both microglia and astrocytes can synthesise chemokines, primary astrocytes are the more evident producer of CCL2, CXCL1 and CXCL10 in response to IL-1β (Fig. 5a). In the APP/PS1 brain, reactive astrocytes proximal to the plaques were not positive for these chemokines in disease *per se* but made higher levels of these chemokines upon local application of IL-1β. This is highly analogous to the exaggerated chemokine responses we recently described in primed astrocytes in the ME7 prion diseased brain (Hennessy et al. 2015) and replication of this phenotype in a distinct disease state, indicates that such priming of astrocytes may, like microglial priming (Perry et al. 2007; Holtman et al. 2015), be a generic feature of astrocytes exposed to some prior neurodegenerative stimulus. Transient hypersensitivty of astrocytes to TLR2 stimulation was previously described, *in vitro*, to occur via prior IL-1β and TNF-α exposure (Henn et al. 2011). More recently astrocytes in the aged brain were also shown to produce heightened expression of some astrocyte-expressed transcripts in response to LPS-induced sepsis (Clarke et al. 2018). Here, we show astrocyte hypersensitivity to IL-1β stimulation after long-term prior exposure to Aβ plaques. Interestingly, in both our studies in ME7 (Hennessy et al. 2015) and in APP/PS1 mice (this study) as well as in the prior *in vitro* study (Henn et al. 2011) and the recent aging study (Clarke et al. 2018), the measure of exaggerated astrocyte responses was increased chemokine output and we showed that this had consequences for neutrophil, monocyte and T-cell infiltration in the ME7 model (Hennessy et al. 2015) and obviously this will require testing in the current model.

Interesting recent studies suggest that astrocytes adopt either A1 or A2 phenotypes depending on *in vitro* innate immune stimulation with IL-1β, TNFα and C1q (Liddelow et al. 2017) and this A1 phenotype is also reported to occur in aged astrocytes (Clarke et al. 2018). However earlier *in vitro* studies using combinatorial stimulation with LPS, TGFβ1 and IFNγ produced multiple divergent astrocyte phenotypes (Hamby et al. 2012). Likewise, *in vivo* stimulation with different pathological stimuli (sepsis vs. medial cerebral artery occlusion) showed significant divergence in astrocyte activation signatures (Zamanian et al. 2012) and astrocytes in normal aging showed significant elevation of many A1 and A2-associated transcripts together. It seems unlikely that astrocytes in the pathological brain could be defined by just 2 phenotypes and the current data, showing phenotypic switching of astrocytes proximal to Aβ plaques, are consistent with the idea of multiple and dynamic astrocyte phenotypes.

We observed hypertrophic GFAP-positive astrocytes around the plaques, distinguishable from astrocytes distal to the plaques, as previously described (Orre et al. 2014). Here, the alteration of a panel of genes chosen from multiple prior astrocyte transcriptomic studies (Zamanian et al. 2012; Orre et al. 2014; Liddelow et al. 2017; Boisvert et al. 2018) showed that transcripts clustered into at least 4 patterns: those not significantly altered by disease or IL-1β: *S100a10*, *Myl9*, *Mertk* (data not shown), those altered by disease: *Gfap* (pan), *Serping1* (A1), *Ctss* and *Il1r1*; those altered by IL-1β but equally in both strains: *Steap4* (pan), *Vim* (pan), *Ccl2, Cxcl1, Ccl5* and *Gper1*; and finally those showing exaggerated responses to IL-1β in Tg mice: *S1pr3* (pan), *Gbp2* (A1), *Clcf1* (A2), *Cxcl10* (pan), *Ptx3* (A2), *Tgm1* (A2), *Cnn1* and *Cst7*. Further characterisation of these profiles will be necessary using isolated cell populations rather than hippocampal homogenates, but the appearance of pan reactive, A1- and A2-associated genes in each of these clusters argues against 2 simple states of activation and cautions against the designation of astrocytes as having an A1 or A2 phenotype based on expression of selected transcripts from that proposed designation.

Moreover the demonstration of exaggerated chemokine expression specifically in the astrocytes proximal to Aβ plaques demonstrates that these astrocytes adopt one phenotype by their proximity to Aβ plaques and adopt another once exposed to local IL-1β. Although enhanced protein expression of CCL2, CXCL1 and CXCL10 has been demonstrated here, only CXCL10 showed exaggerated mRNA expression in Tg animals, suggesting that there may be significant post-transcriptional regulation as well as transcriptional regulation of the astrocyte phenotype.

### Priming vs. training

Although the current study has focussed on the impact of acute inflammation superimposed upon brain already made vulnerable by evolving amyloid pathology, there is also evidence that systemic inflammation in young adulthood (Wendeln et al. 2018), or indeed *in utero* (Krstic et al. 2012), can produce differential responses in glia later, when amyloid pathology is developing. The former has recently been described as innate immune training (Wendeln et al. 2018), based on terminology arising in the peripheral immune literature (Netea 2013). We contend that microglia, and indeed astrocytes, may be trained by multiple different stimuli to produce differential responses to secondary stimuli at a later date and that the established concept of priming (Cunningham et al. 2005) is inclusive of the many potential stimuli that may train microglia for differential responses. The nature, timing and dose of priming stimuli will all impact on microglial and astrocyte phenotypes. The epigenetic underpinning of the training of microglia (Wendeln et al. 2018) provides one potential mechanism underpinning some forms of microglial priming, although with many different stimuli demonstrated to prime microglia (see (Cunningham 2013) for review), there are likely a number of other mechanisms involved. Here we show exaggerated transcription and translation of IL-1 as well as inflammasome assembly in microglia and both transcriptional and post-transcriptional changes in astrocytes, so exaggerated responses of primed glia may be manifest through multiple cellular mechanisms. Mechanisms by which astrocytes become primed to show analagous trained responses now require investigation.

### Systemic inflammation has negative outcomes: from mice to humans

Obviously LPS does not frequently occur within the brain (Singh and Jiang 2004; Chakravarty and Herkenham 2005; Banks and Robinson 2010). However we also show that these microglia change their phenotype when IL-1β arises locally (as is known to occur in situations such as stroke and traumatic brain injury) and also when LPS was administered peripherally as occurs in Gram-negative infection and sepsis. Sepsis leads to intensive care unit admission and high mortality (Iwashyna et al. 2010; Dal-Pizzol et al. 2014) and brain effects are clinically manifest as septic encephalopathy or sepsis-associated delirium (Ebersoldt et al. 2007; Cunningham and MacLullich 2013). Pathophysiological understanding of delirium remains limited and the demonstration here that low dose LPS produces acute deficits selectively in APP/PS1 animals is significant. All animals trained on reference memory in this Y-maze task maintained, despite LPS challenge, their prior performance (ie retention of a well-trained, long-held memory) but when the exit’s location was altered and the animals had to abandon their prior strategy and use visuospatial cues to find a new exit those APP/PS1+LPS animals were worst affected. The impairment of these dynamic cognitive processes simultaneous with preservation of pre-LPS long-term visuospatial memory is typical of the cognitive impairments observed in human delirium, where patients show impairments in cognitive tasks involving processing of novel, trial specific, information but preservation of previously acquired long-term memories (Brown et al. 2011).

It is also significant that LPS-induced sickness behaviour (Dantzer 2004) was equivalent in WT and Tg. One interpretation of exaggerated LPS-induced cognitive deficits occuring selectively in animals with prior diease or old age has been that this is explained simply by the fact that they also show exaggerated sickness behaviour responses (Combrinck et al. 2002; Godbout et al. 2005; Cunningham and MacLullich 2013). That sickness is equivalent in the current study but cognition is disproportionately affected in APP/PS1+LPS mice means that the cognitive impairments are not explained simply as part of the sickness behaviour response and suggests that exaggerated microglial or astrocyte activation, selectively in the circuits underlying these cognitive tasks, or neuronal vulnerability in those circuits can explain the deficits independent of exaggerated sickness behaviour.

The interface of delirium and dementia remains remarkably understudied given that delirium significantly increases the risk of subsequent dementia and/or long-term cognitive decline (Witlox et al. 2010; Davis et al. 2017) and systemic inflammation, a major trigger for delirium, also accelerates cognitive decline in Alzheimer’s disease (Fong et al. 2009; Holmes et al. 2009). Several studies have shown that serial LPS challenge schedules drive increased APP processing, Aβ plaque area and Tau hyperphosphorylation (Kitazawa et al. 2005; Lee et al. 2008; Bhaskar et al. 2010; Wendeln et al. 2018) but recent epidemiological data have shown that patients who receive a diagnosis of dementia after having experienced an episode of delirium showed a weaker association between dementia status and amyoid or Tau (Davis et al. 2012, 2017). Those data suggest that some *de novo* pathology occurs during episodes of delirium and consistent with this we have shown that a single dose of LPS induces new neuronal death (Cunningham et al. 2005) and alters disease trajectory in mice (Cunningham et al. 2009). Therefore, model systems probing these interactions of systemic inflammation provide the possibility to unpick *de novo* pathological events occuring during acute inflammatory episodes. We predict that both astrocyte and microglial priming will play significant roles in systemic inflammation-induced exacerbation of Alzheimer’s disease pathology. Mouse model data show that heightened IL-1 responses drive acute neuronal dysfunction and damage (Skelly et al. 2018) and IL-1 exacerbates Tau pathology (Bhaskar et al. 2010) and “trained” microglial responses influence Aβ levels in the parenchyma (Wendeln et al. 2018). Therefore the potential of targeting the interaction between underlying disease and secondary inflammation is an area of significant therapeutic interest.

However human evidence for microglial and astrocyte priming is lacking. It is now important to identify reliable biomarkers that might demonstrate these concepts in humans. PET imaging of microgliosis can be achieved using 18kDa translocator protin (TSPO) imaging but this has uncertain capacity to distinguish between microglial phenotypes or indeed even to distinguish between microglia and astrocytes (Notter et al. 2018). Cerebrospinal fluid respresents an important opportunity to sample brain levels of cytokines that may inform on local glial phenotypes. To address priming in Alzheimer’s disease, ideally one would be able to sample the CSF of patients with and without established dementia who experience acute inflammatory episodes in order to compare whether those with dementia plus acute inflammatory trauma display exaggerated IL-1β and chemokine concentrations compared to trauma in cognitively normal individuals. These types of studies are beginning to emerge from hip fracture cohorts who receive spinal anaesthesia before replacement surgery (MacLullich et al. 2011; Cape et al. 2014; Skrede et al. 2015) but to address this particular research question such studies require better pre-fracture cognitive and imaging characterisation as well as the obvious need to scale up these very small studies. A growing panel of biomarkers including neurofilament light, neurogranin and synaptic proteins can inform on progression in AD (Hampel et al. 2018) and examining how their levels associate with acutely elevated CSF cytokine/chemokine concentrations is currently being addressed (Ascribed Study Group 2018) and should provide useful points of entry into understanding how acute changes in glial phenotype may contribute to worsening progression in AD.

## Conclusion

We have shown that both microglia and astrocytes are primed by amyloid pathology to show exaggerated IL-1 and chemokine responses to acute inflammatory insults and that these insults have cognitive consequences. It is now a priority to investigate this phenomenon in AD patients and to investigate their contribution to delirium and to progression of dementia.

## Acknowledgements

This study was supported by a Wellcome Trust Senior Research Fellowship to CC (SRF090907) and by NIH R01AG050626. The technical assistance of Gavin MacManus in the Biomedical Sciences Imaging suite is gratefully acknowledged.

